# The FGFR inhibitor Rogaratinib reduces microglia reactivity and synaptic loss in TBI

**DOI:** 10.1101/2024.06.03.597197

**Authors:** Rida Rehman, Albrecht Froehlich, Florian olde Heuvel, Lobna Elsayed, Tobias Boeckers, Markus Huber-Lang, Cristina Morganti-Kossmann, Francesco Roselli

## Abstract

**Background:** Traumatic brain injury (TBI) induces an acute reactive state of microglia, which contribute to secondary injury processes through phagocytic activity and release of cytokines. Several receptor tyrosine kinases (RTK) are activated in microglia upon TBI, and their blockade may reduce the acute inflammation and decrease the secondary loss of neurons; thus RTKs are potential therapeutic targets. We have previously demonstrated that several members of the FGFR family are transiently phosporylated upon TBI; the availability for drug repurposing of FGFR inhibitors makes worthwhile the elucidation of the role of FGFR in the acute phases of the response to TBI and the effect of FGFR inhibition.

**Methods:** A closed, blunt, weight-drop mild TBI protocol was employed. The pan-FGFR inhibitor Rogaratinib was administered to mice 30min after the TBI and daily up to 7 days post injury. Phosphor-RTK Arrays and proteomic antibody arrays were used to determine target engagement and large-scale impact of the FGFR inhibitor. pFGFR1 and pFGFR3 immunostaining were employed for validation. As outcome parameters of the TBI injury immunostainings for NeuN, VGLUT1, VGAT at 7dpi were considered.

**Results:** Inhibition of FGFR during TBI restricted phosphorylation of FGFR1, FGFR3, FGFR4 and ErbB4. Phosphorylation of FGFR1 and FGFR3 during TBI was traced back to Iba1+ microglia. Rogaratinib substantially dowregulated the proteomic signature of the neuroimmunological response to trauma, including the expression of CD40L, CXCR3, CCL4, CCR4, ILR6, MMP3 and OPG. Protracted Rogaratinib treatment exhibited a neuroprotective effect on neuronal density at 7dpi and limited the loss of excitatory (vGLUT+) synapses.

**Conclusion:** The FGFR family is involved in the early induction of reactive microglia in TBI. FGFR inhibition selectively prevented FGFR phosphorylation in the microglia, dampened the overall neuroimmunological response and enhanced the preservation of neuronal and synaptic integrity. Thus, FGFR inhibitors display potential as microglial modulators in TBI.

## Background

Traumatic Brain Injury (TBI) is characterized by the dynamic interplay of multiple cellular actors, including neurons, astrocytes, microglia as well as vascular and immune cells, which may assume beneficial or detrimental roles, depending on time and space (Morganti-Kossmann et al., 2019; Jassam et al., 2017). Microglial cells swiftly react to TBI by migrating to the site of injury (Davalos et al., 2005), assuming an ameboid, chemotactic morphology (Zanier et al., 2015) and diverse reactive functional states including ones characterized by increased interferon response (Somebang et al., 2021) and by disease-associated-like microglial markers (Krasemann et al., 2017; Froehlich et al., 2022). Although unchecked acute microglial reactivity in TBI has been largely considered detrimental, leading to acidosis, oxidative stress, enhanced neuronal damage and synaptic loss (Ritzel et al., 2021; Wang et al., 2017; Jamjoom et al., 2021) in the “secondary injury” phase.

However, early post-traumatic depletion of microglia by CSF1R inhibitor administration reduces the extent of neuronal apoptosis but does not affect the overall lesion size and actually increased the size of intracerebral haematoma (Wang et al., 2022). Actually, a number of microglia-associated responses, such as glial limitans repair and debris clearing may have neuroprotective outcomes (Roth et al., 2014; Russo et al., 2016 ; Loane and Kumar, 2016) and may be carried out by specific subset or functional states of microglial cells (such as repopulating microglia; Willis et al., 2020; Ritzel et al., 2023). Thus, the goal of suppressing microglial reactivity should be substituted by the aim for a fine-tuning microglial reactivity and phenotype to maximize tissue preservation.

Receptor tyrosine kinases have emerged in the last 20 years as a class of drug targets, with more than 70 small-molecule kinase inhibitors approved for human use (mainly in oncology; Roskoski, 2023; Pottiers et al., 2020) and therefore lend themselves to effective drug repurposing. Notably, RKT activation is a prominent response in TBI: a targeted phosphoproteomic screening of 39 RTK has revealed the significant increase in phosphorylation of multiple families of RTK including VEGFR1-3, EphB4, Met, MSPR, EGFR, ErbB3 and FGFR4 at 3h and 24h timepoints (Rehman et al., 2022). In fact, Met and VEGFR, among others, were shown to be phosphorylated in microglial cells, contributing to their regulation. Proof of concept of the use of RTK as entry points for acute TBI treatment has been provided by the use of VEGFR and Met small-molecule inhibitors: both caused the substantial divergence in the phosphoproteomic profile after TBI (demonstrating target engagement) and resulted in improved motor performance (Rehman et al., 2022). Furthermore, prolonged treatment with a Met inhibitor delivered persistent improvement in motor performace and enhanced neuronal preservation (Rehman et al., 2022). These findings have opened up the possibility that multiple RTKs may be involved in the early induction of reactive microglial phenotype(s) and may lend themselves to therapeutic modulation. Among these, some FGFR family members displayed transient up-phosphorylation between 3h and 24h after trauma. Interestingly, the FGFR family does not only control proliferation and plasticity, but has been implicated also in the control of inflammation: blockade of FGFRs reduce the cytokine storm and macrophage proliferation in sepsis (Huang et al., 2019) and FGFR ligand FGF23 induces TNF-α in macrophages (Han et al., 2016). Since FGFR inhibitors have been recently introduced in clinical practice (Weaver et al., 2021), we set out to explore the potential role of FGFR in the early stages of the neuroinflammatory response to TBI.

## Results

### Pan-FGFR inhibitor modifies the RTK phosphorylation landscape of acute TBI

We explored the effect of the pan-FGFR inhibitor BAY 1213802 (Rogaratinib-HCl; Grünewald et al., 2019; henceforth BAY121), on the TBI-associated RTK phosphorylation landscape with the goal to demonstrate effective and specific target engagement. Mice were subjected to a blunt weight-drop mild TBI (or sham surgery), followed 30 min later by administration of BAY121 (or vehicle). NSS score ranged between 0 and 1 (coherently with the mild TBI protocol) at the 3h timepoints. At 3h post injury mice were sacrificed, samples from the injury site were obtained (Fig. 1A) and processed for RTK phosphorylation screening using a nitrocellulose antibody arrays (RTK targets including FGFR 2, 3 and 4). Principal component analysis (PCA) demonstrated a substantial overlap among S-Sal, T-BAY and S-BAY samples, whereas T-Sal stood out (Fig. 1B). Analysis of individual RTK phosphorylation revealed a significant upregulation of p-FGFR3 and p-FGFR4 (but not of pFGFR2) upon TBI which was negated by the BAY121 treatment (Fig. 1C-E). In addition, TBI upregulated the phosphorylation of HGFR, cRet, ErbB4, EphA3 and downregulated phopsho-EphB2 (Fig 1F-J). Interestingly, BAY121 also blocked the phosphorylation of ErbB4 (Fig. 1F) but did not affect the up-phosphorylation of cRet and EphA3 (fig. 1H-I; only statistical trends were detected for HGFR nd EphB2).

**Figure 1.**
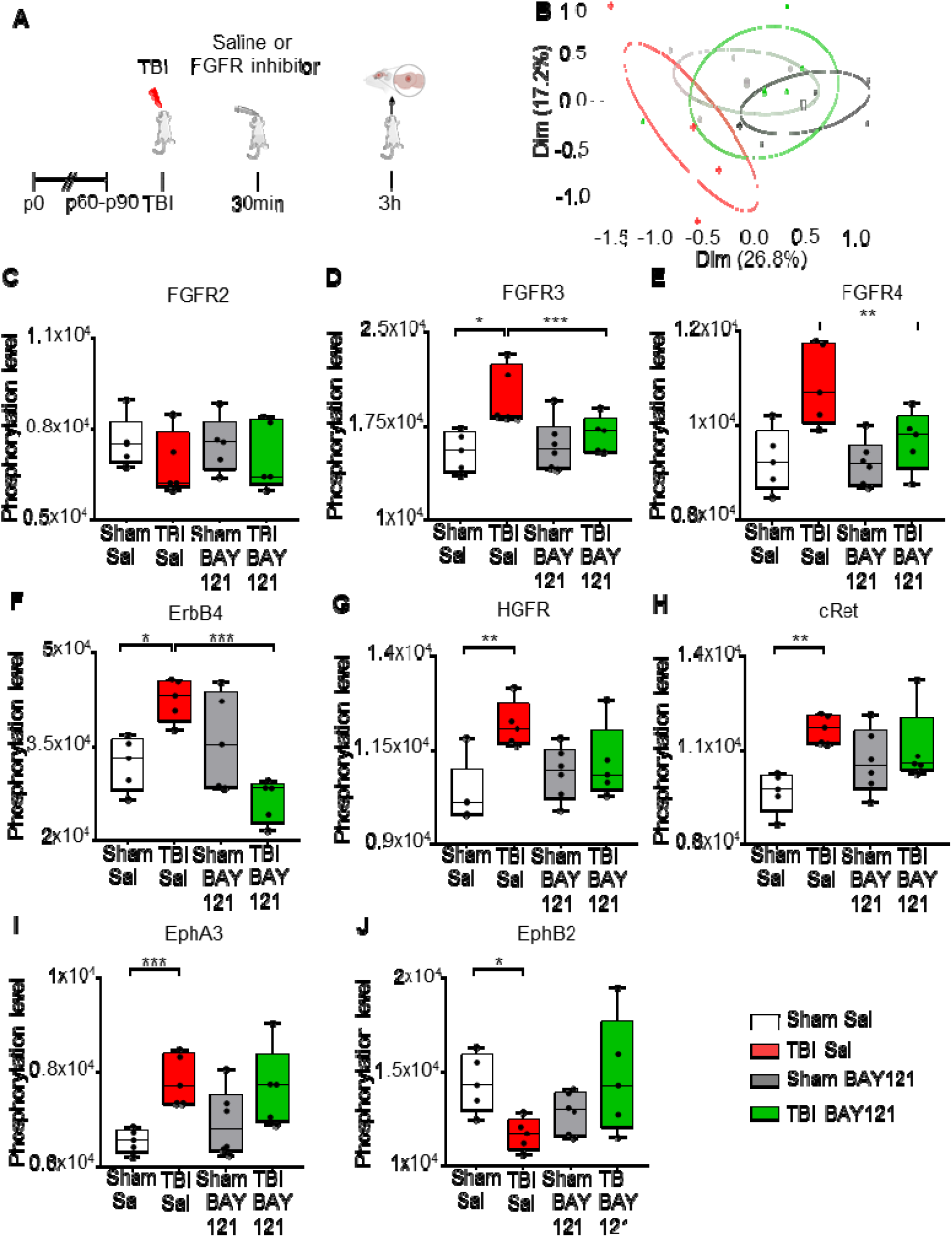
FGFR Inhibitor selectively alters RTK phosphorylation pattern at 3h post injury. (A) Outline of the experimental design. (B) Principal component analysis (PCA) plot displayed group wise distribution of samples. The group specific ellipses indicate 95% confidence interval. (C-J) Differential phosphorylation analysis showed the inhibitor significantly decreased the phosphorylated levels of (D) pFGFR3, (E), pFGFR4 and (F) ErbB4 after TBI. However, the inhibitor showed no effect on phosphorylation levels of (C) FGFR2, (G) HGFR, (H) cRET, (I) EphA3, and (J) EphB2. Significance for differentially phosphorylated proteins was set at p<0.05 (FDR adjusted). (B-J: n=5-6/group; ns= not significant, *p<0.05, **p<0.01, ***p<0.001).

Taken together, these findings demonstrate that TBI upregulates the phosphorylation of FGFR and the BAY121 FGFR inhibitor successfully negates this event; BAY121 (confirming target engagement) does not appear to block the phosphorylation of other RKTs with the exception of ErbB4 (supporting the selectivity of BAY121).

### FGFR inhibitor prevents FGFR1 and FGFR2 phophorylation in microglia upon trauma

We investigated the cellular sources and the spatial distribution of FGFR phosphorylation using immunohistological approaches; antibodies against FGFR1(pY654) and FGFR3(pY724) proved suitable for immunolabeling of brain sections. Animals were subject to TBI and injected 30 min later with either saline or BAY121 and sacrificed 3h after the injury (2.5h after treatment; figure 2A). At this timepoint, immunoreactivity for both pFGFR1 and pFGFR3 was significantly upregulated in the site of injury in T-Sal but not in T-BAY samples (Fig.2B, E). Interestingly, the immunoreactivity pattern of pFGFR1 and 3 highlighted large number of small cells of ramified morphology, resembling microglia. In fact, co-immunostaining with Iba1 demonstrated that >90% of Iba1+ cells in the site of injury displayed immunoreactivity for pFGFR1 and >50% of Iba1+ cells displayed immunoreactivity for FGFR3. In T-Sal samples, both the immunofluorescence intensity in Iba1+ cells (Fig. 2C, F) and the fraction of Iba1+ immunopositive for phospho FGFR1 or phospho FGFR3 was significantly increased compared to S-Sal (Fig. 2C-D and 2F-G). Notably, in T-BAY samples, both the immunofluorescence intensity and the number of Iba1+ cells immunopositive for phosphoFGFR1 and FGFR3 were strongly decreased (no change in the total number of Iba1+ cells was noted).

**Figure 2.**
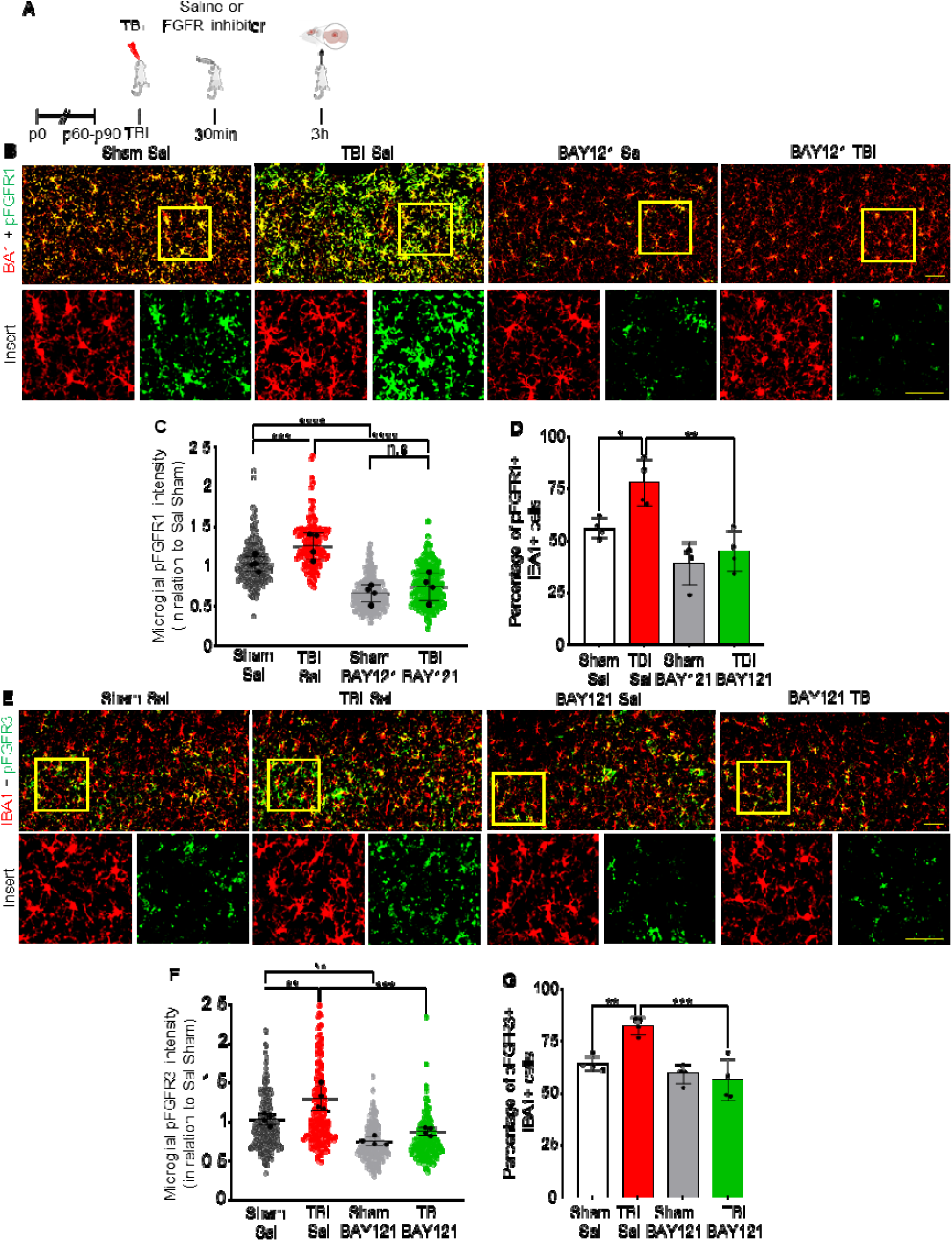
Upregulation of pFGFR1 and pFGFR3 in microglia 3h post injury. (A) Outline of the experimental design. (B) Immunostainings for pFGFR1 (cyan) and IBA1 (magenta) in the left column together with pFGFR3 (cyan) and IBA1 (magenta) in the right column for Sham Sal, TBI Sal, Sham BAY121 and TBI BAY121 treated mice. Scale bar: 50µm. (C) FGFR1 upregulation after TBI is controlled by BAY121 treatment. (n=4/group; n.s. = not significant; ***p<0.001; ****p<0.0001) (D) Immunostaining for pFGFR3 shows it is upregulated in microglia but this effect is negated by BAY121. (n=4/group; n.s. = not significant; **p<0.01; ***p<0.001)

Taken together these results confirm the elevation in FGFRs activation upon TBI and demonstrated the successful target engagement for BAY121 on microglial cells.

### FGFR inhibition significantly suppresses immune responses in the site of injury

Next we explored if prolonged BAY121 administration could not only affect the acute reactivr microglial phenotype, but generate a long-lasting, broad alteration of the proteomic neuroimmunological landscape associated with brain injury. Mice subjected to trauma were administered with BAY121 (or vehicle) 30 mins after trauma (or sham surgery) and continued daily for 3 days (Fig. 3A). A targeted proteomic profile of the injury site, involving >1300 individual protein was obtained by antibody arrays. PCA plot showed a substantial separation between S-Sal and TBI-Sal samples on one side and S-BAY and T-BAY on the other (Fig. 3B). When compared to S-Sal samples, TBI-Sal samples displayed the upregulation of 16 proteins and the downregulation of 3 proteins (Fig. 3C). The upregulated proteins involved multiple mediators of inflammatory responses, including KC (murine homologue of the chemoattractant IL-8), the microglial regulator Axl, the chemotactic receptor CCR10 and the immune regulator CD40 and its ligand CD40L. On the other hand, BAY121 resulted in a substantial change in the proteome when administered after trauma: 51/1308 proteins were downregulated and 98/1308 were upregulated in TBI-BAY121 vs TBI-Sal samples (Fig. 4D-E, Supplementary Fig. 1A). The gene ontology analysis revealed that BAY121 treatment resulted in the downregulation of proteins involved in immune function and cytokine response (top 5 GO categories, Supplementary Fig. 1B-C). Strikingly, Axl and CD40L (upregulated in TBI-Sal vs Sham-Sal) were among the top strongly downregulated along with microglia polarization indicator CD80 and, proinflammatory signaling and chemotaxis markers such as CXCR3, CCL4, CCR4, ILR6, MMP3 and Osteoprotegerin (recently involved in microglial reactivity; Froehlich et al., 2022) (Fig. 3D). The upregulated proteins display an enrichment in the GO categories of cellular signaling, metabolism and protein synthesis, including the mitochondrial protein TRAP and the ion channel KCNAb3.

**Figure 3.**
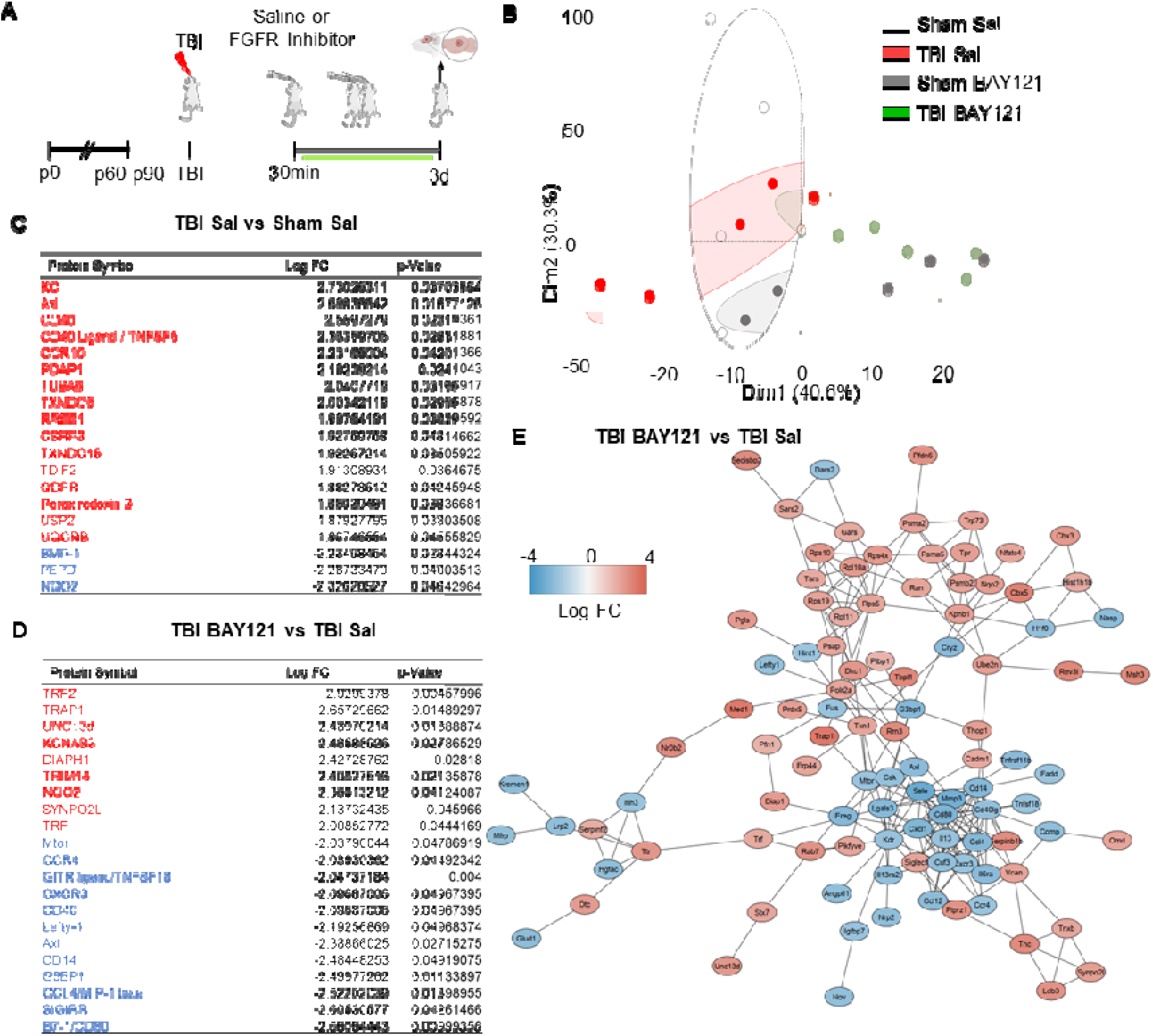
FGFR inhibitor suppresses immune-related proteome signature at 3d post injury. (A) Outline of the experimental design; BAY121 was administered 30 mins after trauma and continued for 3d (1 dose/day). Samples were collected 3d post injury. (B) PCA plot showed distribution of groups across two dimensions contributing to >70% variability among the groups. (C-D) After modified DP analysis (FDR <0.05), subset of significantly upregulated (Red) and downregulated (Blue) proteins with log fold change and individual significance for (D) TBI-Sal compared to Sham-Sal and (D) TBI-BAY121 compared to TBI-Sal (n=5/group). (E) Protein-protein-interaction analysis revealed distinct clustering of downregulated and upregulated proteins in TBI-BAY121 vs TBI Sal; the cluster of downregulated proteins is enriched with immune- and inflammation-related proteins.

**Figure 4.**
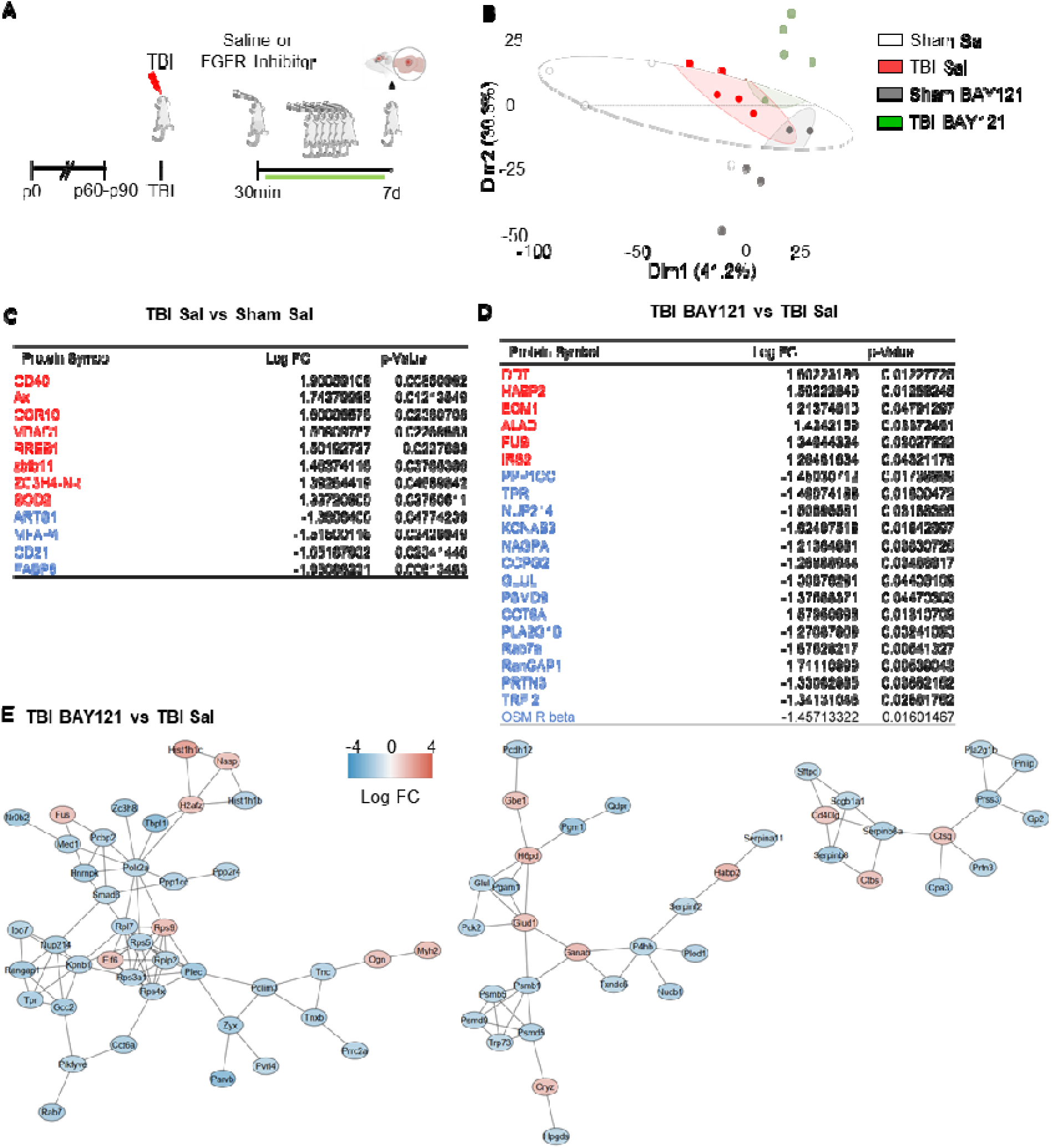
Prolonged FGFR inhibitor treatment significantly alters the TBI-related proteome profiles at 7d post injury. (A) Outline of the experimental design; BAY121 was administered 30 mins after trauma and continued for 7d (1 dose/day). Samples were collected 7d post injury. (B) PCA plot showed distribution of groups across two dimensions contributing to >70% variability among the groups. (C-D) After modified DP analysis (FDR <0.05), subset of significantly upregulated (Red) and downregulated (Blue) proteins with log fold change and individual significance for (C) TBI-Sal vs Sham Sal and (D) TBI-BAY121 vs TBI-Sal. (E) Protein-protein interaction analysis revealed three distinct networks affected by prolonged BAY121 treatment, related to protein synthesis proteasomal degradation and inflammation (n=5/group)

We further characterized the distinct signature imposed by BAY121 treatment upon TBI by constructing a protein-protein interaction (PPI) network for the proteins up- or down-regulated by the treatment. We employed the String algorithm and visualized the network using the Cytoscape software. After preprocessing of the dataset, PPI network displayed 137 nodes and 243 edges (Fig. 3E). Most notably, proteins up- and down-regulated by BAY121 were enriched in two distinct clusters, with most of the down-regulated proteins related to the immune response forming a tight cluster. Taken together, the proteomic data suggest that prolonged administration of the FGFR inhibitor BAY121 profoundly reduces the sub-acute neuroinflammatory response to TBI.

### Prolonged FGFR inhibitor alters both neuron-specific and immune-specific proteins 7d post injury

We further investigated the proteomic signature of prolonged FGFR inhibition in TBI by taking into consideration samples obtained at 7 dpi. As before, mice subjected to trauma were administered with BAY121 (or saline vehicle) 30 mins after trauma. The treatment was continued for 7 days (1 dose/day) for 7 days (Fig. 4A). PCA plot based on the targeted proteomics (1308 targets), revealed that, while the S-Sal and T-Sal samples largely clustered together, S-BAY121 and TBI-BAY121 minimally overlapped the Sal groups (Fig. 4B), indicating a persistent and profound effect of BAY121 treatment.

The analysis of differentially expressed proteins revealed the increased expression in TBI-Sal (vs Sham-Sal) of the proteins involved in inflammation such as Axl, CD40, CCR10 and SOD2, whereas the B-cell marker CD21 was downregulated (Fig. 4C). Notably, the comparison of BAY121-treated TBI vs vehicle-treated TBI samples revealed a substantial divergence in the proteome: 123/1308 proteins were downregulated and 38/1308 proteins were upregulated (Fig. 4D, E). The gene ontology analysis of the downregulated proteins revealed a substantial involvement of proteasome regulatory proteins and nucleo-cytoplasmic trafficking, pointing toward an impaired protein degradation and cellular stress (Supplementary Fig. 2B). The comparatively small number of upregulated proteins did not lend itself to a reliable GO analysis. Interestingly, when we mapped the PPI of proteins altered by BAY121 treatment (TBI-BAY121 vs TBI-Sal) using STRING/Cytoscape, three distinct sub-networks (cumulatively displaying 112 nodes and 135 edges; Fig. 5E) were identified. The smallest of the network (13 proteins) involved downregulated proteins related to the inflammatory/phagocytic function (notably Catepsins and Serpins) and immune regulation (such as CD40L). The two larger networks (65 proteins) included many subunits of the proteasome system, chaperones and trafficking proteins, the largest majority of which were downregulated (Fig. 4E, Supplementary Fig. 2A). Based on these findings, the proteomic analysis at 7dpi suggested that blockade of FGFR signaling produced a persistend impact on the neuroinflammatory cascade but also substantially impacted the tissue protein homeostasis.

**Figure 5.**
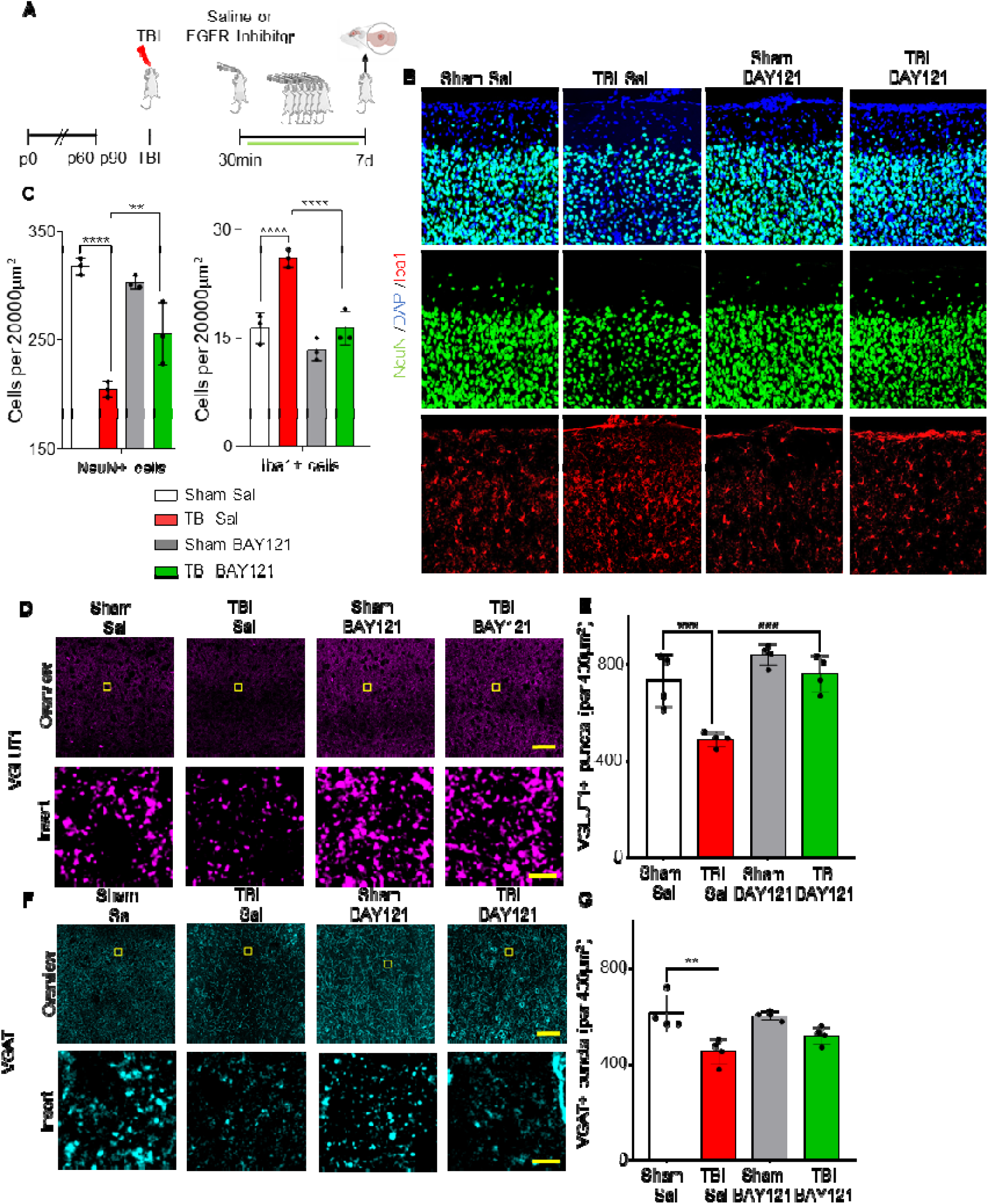
Prolonged FGFR inhibitor administration preserves neuronal density at 7 dpi. (A) Outline of the experimental design; BAY121 was administered 30 mins after trauma and continued for 7d (1 dose/day). Samples were collected 7d post injury. (B-C) Immunostaining for NeuN showed a significantly decreased number of neurons in the injury site of TBI-Sal group compared to Sham-Sal; a significantly higher number of neurons was seen in the TBI-BAY121 group. Note that microglial density was also increased in TBI-Sal but not in TBI-BAY121. (n=4/group; **p<0.01, ***p<0.001, ****p<0.0001) (A) Loss of VGLUT1 density after TBI is dependent on FGFR signaling 7d after TBI. (n=4/group; ***p<0.001; Scale bar: overview = 70µm; insert = 5µm) (C) VGAT density in the TBI core is significantly reduced at 7dpi. (n=4/group; **p<0.01; Scale bar: overview = 70µm; insert = 5µm)

### FGFR inhibitor reduces neuronal loss and synaptic loss 7d post trauma

Finally, we sought to determine if the effects of FGFR blockade on the neuroinflammatory response to TBI were associated with reduced neuronal vulnerability and synaptic integrity. As before, mice were subjected to trauma and treated with either saline or the FGFR inhibitor 30 mins after trauma for 7 days (1 dose/day) (Fig. 5A). As expected, density of microglia was still increased in the injury site of TBI -Sal mice, but not in TBI-BAY121 group (Fig. 5B, C). Conversely, the density of NeuN+ cells was significantly reduced in the site of injury in TBI Sal mice compared to Sham mice (Fig. 5B, C). Notably, BAY121 treatment significant increased the number of surviving neurons in the injury site.

We further explored the preservation of synaptic structures in the injury site upon BAY121 treatment. We assessed the density (number of synaptic puncta per 400µm^2^) of excitatory synapses in the form of pre-synaptic VGLUT1 as well as of inhibitory synapses in the form of pre-synaptic VGAT. VGLUT1 showed a significant loss in synaptic density after TBI, which was not observable any more after BAY121 treatment 7dpi (Fig. 5D-E). VGAT+ terminals were also significantly decreased in the lesion area at 7 dpi (Fig. 5F-G) with a trend toward better preservation in T-BAY samples.

## Discussion

Our data show that FGFR family is prominently activated in the early phases of mild TBI in particular in microglial cells. Inhibition of the FGFR family by BAY121 results in the suppression of early microglial reactivity and reduces the neuroinflammatory footprint at later stages, which ultimately leads to an improved preservation of neuronal and synaptic integrity in the site of injury.

TBI results in the simultaneous activation of multiple RTK families, often with distinct temporal dinamics (Rehman et al., 2022): the activation of HGFR/Met has been shown to lead to a reactive, phagocytic microglial state with detrimental consequences (Rehman et al., 2022), whereas the activation of the ErbB family in inhibitory interneurons controls synaptic plasticity and circuit activity after TBI (Chandrasekar et al., 2018). Likewise, activation of the VEGF-C/VEGFR3 contributes to drive microglial polarization after TBI (Ju et al., 2019) and multiple members of the Tyro-Axl-Mer RTK family regulate microglial reactivity and phagocytic activity in TBI (Wu et al., 2021) and other disease conditions (Zhou et al., 2023). Our focused screening, together with the immunohistochemical confirmation, reproduce the activation of Met, ErbB4, EphB2 and ErbB4 previously identified (Rehman et al., 2022; Chandrasekar et al., 2018) and identify a significant phosphorylation of several members of the FGFR family. The pan FGFR inhibitor BAY121 effectively suppresses FGFR1, FGFR3 and FGFR4, with limited effect on the trauma-related activation of other RTK. A notable exception is the full blockade of the induction of ErbB4 phosphorylation; since ErbB4 is highly involved in synaptic plasticity and stability (Bernard et al., 2022; Luo et al., 2021) and it is activated by TBI (Chandrasekar et al., 2018), it is conceivable that the reduced activation may be a consequence of the modulated microglial reactivity and synaptic preservation rather than a direct inhibition. The combined screening and immunohistochemistry data demonstrate that BAY121 is effective and relatively selective in preventing FGFR family receptors activation in microglia when administered 30 min after trauma, i.e. in a therapeutic window.

The blockade of FGFR signaling by BAY121, both in acute (single dose) or protracted (3 to 7 days) administration, consistently results in the dampening of the neuroinflammatory response to TBI at the level of microglial reactivity as well as in terms of broad inflammatory proteomic footprint. Interestingly, FGFR family members are expressed not on a number of immune cells, suggesting their broad contribution to immunity and inflammation: FGFR2 mediates chemotaxis in neutrophils (Haddad et al., 2011), and FGFR1 is expressed in T-cells (Byrd et al., 2003; Fahranak et al., 2017) as well as on macrophages and lymphocytes in lupus nephritis (Rossini et al., 2005). At brain level, FGFR signaling contributes to microglial reactivity to bacterial products by inducing pro-inflammatory cytokines (Parthasarathy et al., 2023) and a small-molecule FGFR inhibitor reduces the infiltration of lymphocytes and activated macrophages, as well as the production of pro-inflammatory cytokines, in the EAE model (Rajendran et al., 2023). Thus, our data are coherent with a role of the FGFR family in modulating the overall neuroinflammatory cascade and cytokine response to injury, including microglial reactivity. Interestingly, FGFR expressed on oligodendrocytes also contribute to the regulation of neuroinflammatory cascades in EAE: conditional deletion of FGFR1 or FGFR2 from oligodendrocytes results in decreased microglial reactivity and lymphocytes infiltration in EAE as well as in reduced levels of pro-inflammatory cytokines (Rajendran et al., 2018; Kamali et al., 2021); thus, FGFR blockade may immunomodulatory effects through additional cell types. Although in our model blockade of FGFR resulted in reduced neuroinflammation and enhanced synaptic and neuronal preservation in our model, anti-inflammatory effects of FGFR have been also reported: systemic administration of FGF21 reduces the inflammatory response in a stroke model (Wang et al., 2020) and intranasal administratino of FGF20 reduced blood-brain-barrier impairment in severe TBI (Chen et al., 2021). Thus, the net effect of FGFR activation may depend on the complex interplay of the inflammatory context and of the mix of FGFR ligands available (Parthasarathy et al., 2023). To this respect, multiple RTK are activated simultaneously in mild TBI (Figure 1 and Rehman et al., 2022); blockade of FGFR does not affect other RTK also involved in regulation of microglial reactivity (such as HGFR/Met), although a similar anti-inflmmatory effect is observed. This may imply a functional redundance of RTK regulation of reactive microglia, or may suggest that the ultimate phenotype is dependent on the combinatorial activation or one or more RTK, which would enable the fine-tuning of the response to the specific conditions. Hence, FGFR signaling may be ultimately pro- or anti-inflammatory depending on the ensemble of RTK activation and the context conditions.

Histological readouts demonstrate that protracted BAY121 administration results in a reduced loss of excitatory and inhibitory synapses and in the overall preservation of neuronal survival. The proteomic analysis highlights the lack of inflammatory mediators in BAY121-treated samples (still present in the T-Sal samples) and the downregulation of two clusters of proteins involved in proteasome regulation, Golgi function, ribosome biosynthesis and a third cluster of proteases and protease inhibitors. The reduced microglial reactivity (reduced DAM-like phenotype and decreased CD68 expression) may contribute to the preservation of synaptic integrity, since microglia mediates synaptic elimination in several disease settings (Cangalaya et al., 2023; Tzioras et al., 2023); the downregulation in multiple proteases and protease inhibitors enriched in phagocytes, such as Cathepsins and Serpins, is compatible with this model. In addition, the downregulation of proteasome-regulators and in trafficking proteins may point toward a more limited synaptic de-stabilization and reduced degradation of synaptic components (many of which, are proteasome-dependent: Ehlers, 2003; Kerrisk et al., 2018; Roselli et al., 2011). It must be stressed that FGF/FGFR themselves are regulator of synaptogenesis and synaptic stability (Li et al., 2002; Daborwski et al., 2015) and, although we observed a positive impact on synaptic preservation, this outcome may be the net consequence of effects on microglial as well as on other cells types.

A few limitations of the present work are worth addressing. BAY121 is a small-molecule pan FGFR inhibitor; however, the possibility of small-molecule tyrosine kinase inhibitors having additional targets (Reinecke et al., 2023; Grünewald et al. 2019), possibly contributing to their biological effect, cannot be fully discounted; to date, our array RTK screening does not demonstrate substantial inhibition besides the FGFR family. Furthermore, the effect of systemic administration of a pan-FGFR inhibitor may be not restricted only to microglia but may involve additional players in the CNS (e.g., neurons, oligodendrocytes, immune cells; Daborwski et al., 2015; Klimaschewski, et al., 2021; Rajendran et al., 2021) although at least in the acute phase, phosphorylation of FGFR1 and 3 is largely restricted to Iba1+ cells. Finally, FGFR are endowed with highly pleiotropic functions in synaptic stability (Daborwski et al., 2015), axonal guidance (Yang et al., 2018) and myelination (Furusho et al., 2015) and the full spectrum of the beneficial and detrimental consequences of FGFR inhibition in acute TBI are not fully elucidated.

## Conclusion

Our findings provide a proof-of-concept of the translational value of targeting the FGFR in acute TBI for the modulation of early microglial reactivity and enhanced preservation of neuronal and synaptic integrity. Recently, rogaratinib-HCl entered clinical trials (Sternberg et al., 2023; Addeo et al., 2022) and three FGFR inhibitors (pemigatinib, futibatinib, and infigratinib) have been approved for human use in the therapy of several gastrointestinal, urologic or haematopoietic neoplasms (Weaver et al., 2021; Subbia et al., 2023). Although their chronic administration is not devoid of side effects (Subbia et al., 2023), our findings support the investigation of their repurposing for short-term modulation of neuroinflammation in TBI.

## Materials and Methods

### Animals

All experimental procedures were performed in compliance with animal protocols approved by the local veterinary and animal experimentation committee at University Ulm and by the Regierungspraesidium Tubingen under the license no. 1370. B6SJL male mice aged between p60–p90 days were used throughout the study.

### Traumatic brain injury procedure

Modified closed, blunt weight drop model Traumatic Brain Injury (TBI) was performed as previously reported (Rehman et al., 2022). For all procedures, mice were anesthetized with sevoflurane (2-4% in 96% O2) and were subcutaneously injected with buprenorphine (0.1mg/kg; 1 dose/day) as a pre- and postoperative analgesic. The scalp was shaved and eye ointment was applied preoperatively to protect the cornea. Scalp skin was then incised on the midline to expose the skull and the animals were positioned in the weight-drop apparatus in which the head was secured to a holding frame. Using the 3-axis mobile platform in the apparatus, the impactor was positioned to the coordinates of the injection site (From bregma ≈ x = +3.0mm, y = − 2.0mm, z = 0.0mm). TBI was delivered by dropping a weight of 120g from a height of 45 cm. A mechanical stop prevented a skull displacement (by the impactor) larger than 2.5 mm, in order to keep the brain damage comparable. Apnea time was monitored after injury. The Neurological Severity Score (NSS) was assessed after 3h, 1 dpi and at 7 dpi and never exceeded the score 1 for any mouse. As such, no animal met the criteria for early sacrifice. Mice were checked every 2 hours on the day of trauma. Effort was made to minimize the suffering of animals and reduce the number of animals used. The contralateral hemisphere was used as control samples throughout the study.

### Neurological Severity Score (NSS) measurement

Throughout all animal experiments the NSS (Flierl et al., 2009) was measured at 3h, 1h and 7d, depending on sacrificial time of the individual mouse. The NSS is comprised of a total of 10 individual tasks mice were subjugated to each timepoint the NSS was measured with a 10 to 30 seconds break inbetween each individual test. The tests include an arena escape within 3min, mono-/hemiparesis, straight walking, search behaviour, startle reflex, balancing on a) a 7mm wide angular beam and b) a 5mm wide round beam and finally a beam walk test with a lenght of 30cm and a with of a) 3cm, b) 2cm or c) 1cm. Points were awarded when mice could not fulfill an individual task, which then were summed up into the total NSS score. The total NSS score for all animals sacrificed as part of the publication are reported in Supplementary Table 1.

### Immunohistochemistry

Brain samples were processed as previously described (Froehlich et al., 2022; Rehman et al., 2022). Briefly, mice were sacrificed by trans-cardial perfusion with 4% PFA in PBS, and brains were dissected and postfixed in 4% PFA overnight. Brains were then transferred to 30% Sucrose for 2 days, after which the samples were embedded in OCT (Tissue Tek, Sakura, Germany). 40 micron sections were cut with a cryostat (Leica CM 1950 AG Protect cryostat). Sections spanning the injury site were selected and blocked (3% BSA, 0.3% Triton in 1x PBS) for 2h at room temperature, followed by incubation for 48h to 72h, depending on Antibody sensitivity at 4°C with primary antibodies diluted in blocking buffer. Sections were washed 3x 30 min with PBS and incubated for 2h at RT with secondary antibodies diluted in blocking buffer. The sections were washed with PBS and mounted using Prolong Gold Antifade Mounting Medium (Invitrogen, Germany). A list of the antibodies used in this study can be found in Supplementary Table 2.

### Image Acquisition and analysis

Confocal images were acquired in 1024 x 1024 pixel and 12-bit format, with a Leica DMi8 inverted microscope, equipped with an ACS APO 40x oil objective. Parameters were set to obtain the optimal signals from the stained antibody or mRNA and at the same time avoiding saturation. All fluorescent channels were acquired independently, to avoid cross-bleed. 3-4 sections spanning the core and perilesional area of the impact site were imaged of each mouse. For each image a tile scan was set up consisting of x by x tiles spanning the injury location.

For image analysis, stacks were collapsed in maximum intensity projection pictures and mean gray value or cell density per fixed region of interest (ROI) was measured. For quantification, we considered a 200µm x 200µm ROI centered on the axis of the injury site.

Synaptic density was detected after producing a mosaic image corresponding to 6 x 6 single optical sections (acquired with a 63x oil objective) with 1 μm thickness. Each cortical section was imaged at a fixed depth 5-10µm inside the section, the composite image was positioned so that an uninterrupted coverage of the impact site with perilesional area was acquired. Quantification of density of synapses was done with the IMARIS software (Bitplane AG, Zurich, CH), ROI (450µm x 450µm) were positioned at fixed distance into the cerebral cortex (layer II/III) at the core of the injury. The number of Vglut and VGAT synapses per unit area was counted using IMARIS. The parameters were kept constant for each ROI.

### Phospho RTK array processing

Proteome Profiler Mouse Phospho-RTK Array Kit (R&D Systems, Minneapolis) was used to determine the Phospho RTK activation pattern. The nitrocellulose membrane arrays provided in the kit were based on sandwich immunoassay and processed according to manufacturer’s instructions. Briefly, membranes spotted with the anti-RTK antibody were blocked in Array buffer 1 for 1h at RT. 160 ug of protein extracted from cortical samples was diluted in 1.5mL Array buffer 1 overnight at 4°C. After washing, arrays were incubated for 2 hours at RT with Anti-Phospho-Tyrosine-HRP Detection Antibody, diluted to 1:5000 in 1X Array Buffer 2. After final washing steps, HRP detection was performed by adding 1 ml Clarity Max™ Western ECL Blotting Substrates from Bio-Rad. Arrays were imaged using BioRad X-ray imager and quantified using ImageJ. ROI was drawn on each antibody spot with a constant diameter and mean gray value was recorded. Further analysis was performed using R software.

### Proteome Array processing

200 ug of protein extracted from cortical samples was loaded to the arrays. The arrays were processed according to the manufacturer’s instructions. Briefly, the sample was biotinylated and was mixed with reaction stop reagent and incubated for 30 mins at RT. The glass arrays were incubated in a blocking solution for 45 minutes and washed. After adding a biotinylated sample to the coupling solution, arrays were incubated for 2 hours at RT and washed. The arrays were incubated with a detection buffer with Cy5-streptavidin (ThermoFisher) at 1:1000 for 20 minutes at RT. The washing steps were repeated as described in the manufacturer’s instructions. After removing excess ddH20 from the slides, the arrays were dried. The arrays were imaged using a GenePix 4000B array scanner (Molecular Devices, LLC) and the image analysis was performed using GenePix Pro Software v7 (Molecular Devices, LLC). The settings for the analysis were kept constant in all cases. The GAL file was loaded in the software and the ROIs were adjusted on the protein spots. Each intensity on F635 was recorded and GPR files were saved. Further analysis was performed using R software.

### Protein-Protein Interaction network and hub genes

We constructed PPI networks to analyze the functional interactions among differentially expressed proteins, using Search Tool for the Retrieval of Interacting Genes/Proteins (STRING: http://www.stringdb.org) (Franceschini et al., 2013) and visualized the networks using Cytoscape (https://cytoscape.org/) (Kohl et al., 2011).

### Array bioinformatic analysis

Chemiluminescence signal for each spot was logged after microarray image analysis. The raw intensity values for each receptor/protein were recorded automatedly via image recorder software. The raw data files were loaded in R software and the dataset for each array was preliminary subjected to quality control assessment (QCA), outlier identification, data distribution, intra-array and inter-array normalization. Normalized data for each array was subjected to principal component analysis (PCA) to display group-based clustering. Confidence ellipses (assuming multivariate normal distribution) with the first two principal components were plotted to validate further analysis. Modified linear modeling-based analysis was then applied to the data to identify significant increase or decrease in phosphorylation or protein level. For protein array analyses, the code has been made publicly available on open-access GitHub repository PROTEAS (Rehman, 2020).

### Statistics

Statistical analysis was performed with the GraphPad Prism software suite. Mann-Whitney U-Test was used for two group comparisons, One-way ANOVA with Bonferroni Correction was performed among three groups and Two-way ANOVA with Tukey correction was used for four group comparisons to examine statistical significance. Protein array analysis was performed using R software. Error bars represent standard deviation (SD), unless indicated otherwise. Statistical significance was set at P < 0.05.

## Supporting information

supplementary table 1-NSS scores

## Declarations

### Ethics approval and consent to participate

All experimental procedures were performed in compliance with animal protocols approved by the local veterinary and animal experimentation committee at University Ulm and by the Regierungspraesidium Tubingen under the license no. 1370

### Consent for publication

NA

### Availability of data and materials

Raw microscopy images datasets and array proteomics and phosphoproteomics dataset are available upon request.

### Author’s contributions

FR, RR and AF designed and supervised the project and wrote the initial manuscript draft and prepared the figures. RR and FoH performed the trauma surgery and collected the samples. RR and LE performed the array proteomics and array phosphoproteomics and the bioinformatic analyis. TB provided antibodies and the use of the confocal microscope. RR, FoH and AF performed the confocal microscopy and image analysis. FR, RR, MHL, CMK, TB and AF contributed to data analysis, the initial and final manuscript drafts and to the preparation of figures.

## Acknowledgements

RR and AF wish to thank the Hannelore Kohl Foundation for its support. All authors wish to thank Thomas Lenk and Gizem Yartas for their dedicated technical support and dr. Diana Wiesner and dr. Jelena Scekic-Zahirovic for discussions and technical suggestions.

## Supplementary figures

**Supplementary figure 1.**
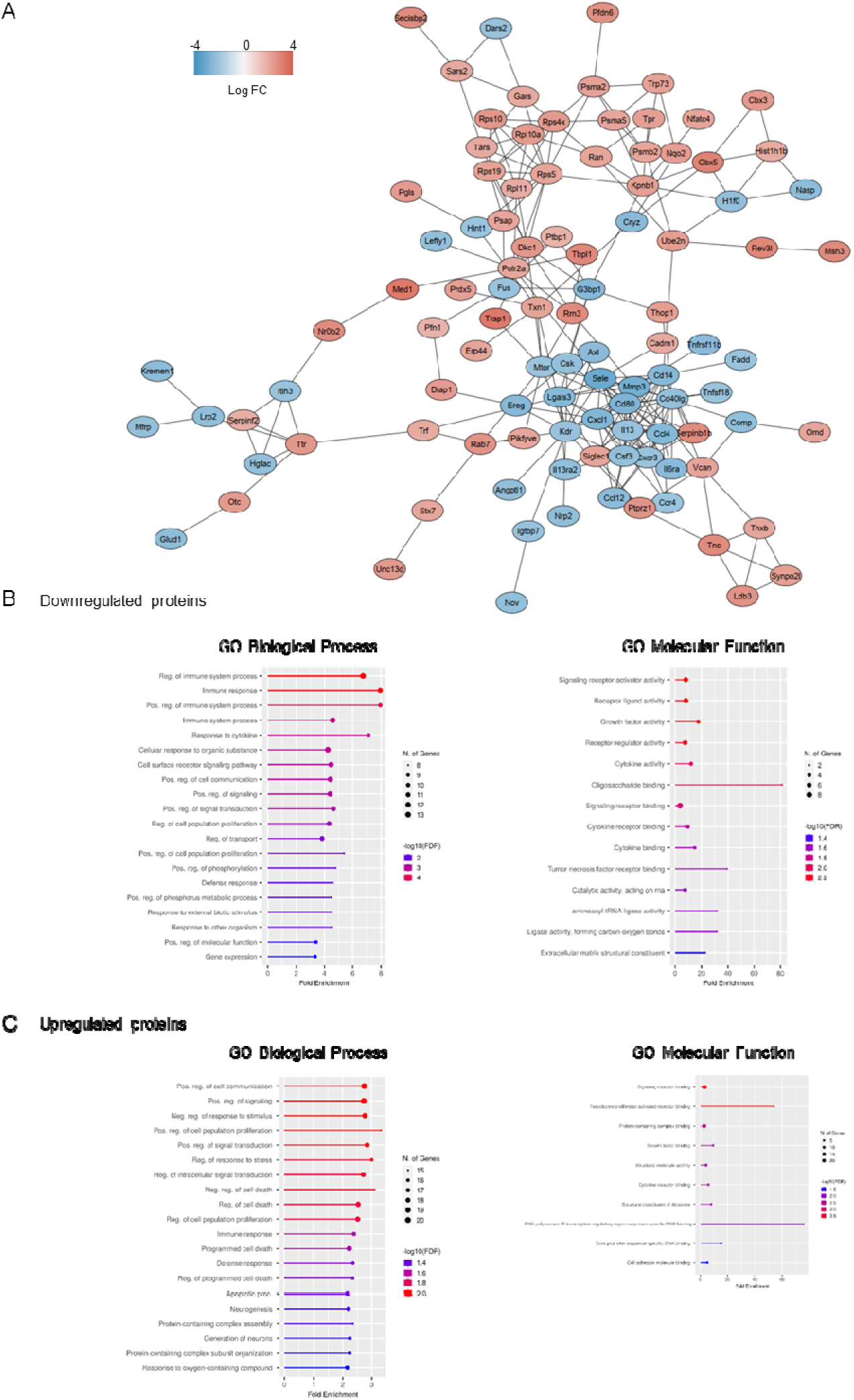
Comparison of TBI sal and TBI BAY121 at 3d post injury. (A) Protein-protein interaction (PPI) of significantly upregulated (red) and down regulated (blue) proteins. Color distribution is based on fold change (FC). (B-C) Gene ontology (GO) analysis and for (B) down regulated proteins and (C) upregulated proteins.

**Supplementary figure 2.**
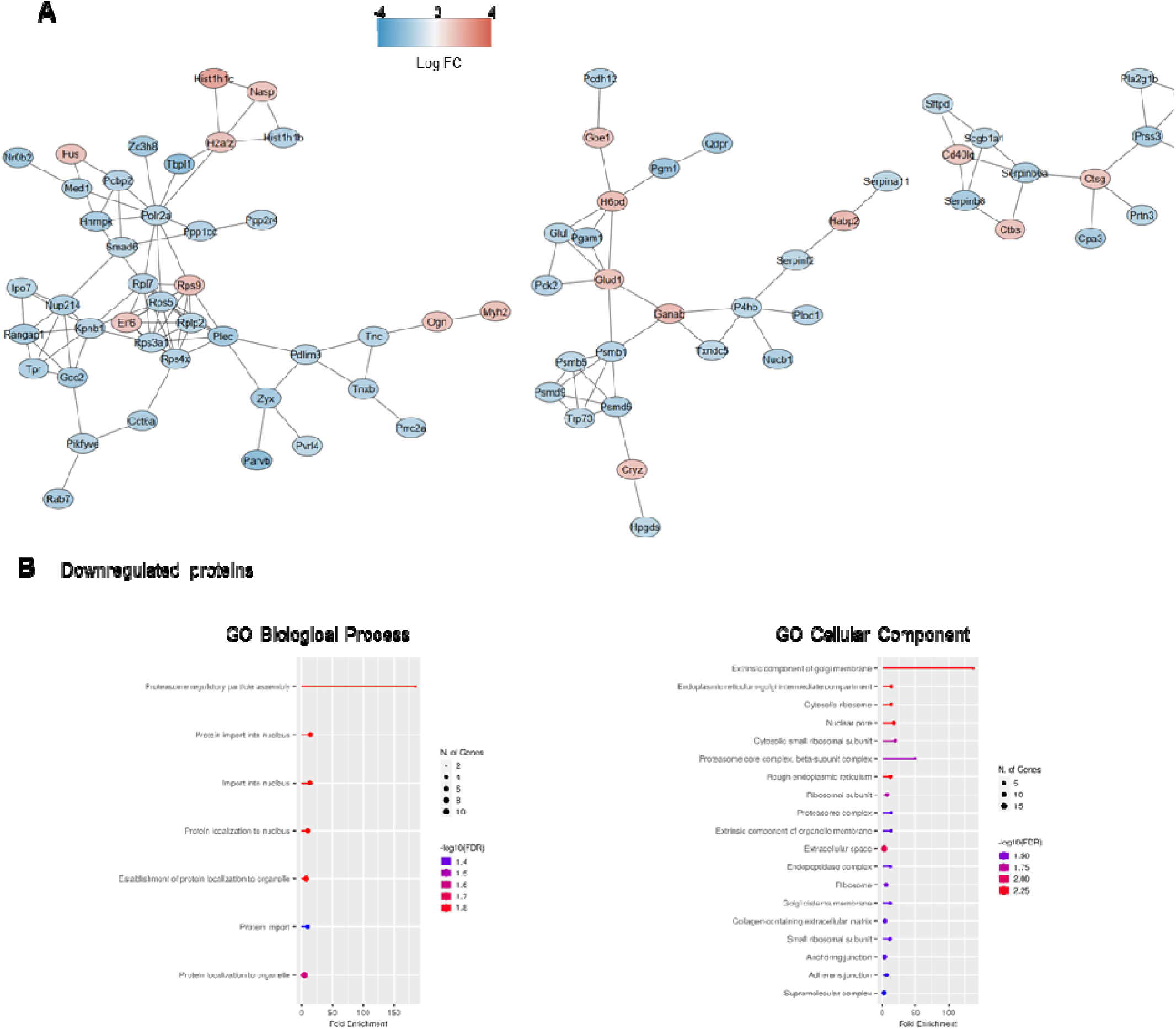
Comparison of TBI sal and TBI BAY121 at 7d post injury. (A) Protein-protein interaction (PPI) of significantly upregulated (red) and down regulated (blue) proteins. Color distribution is based on fold change (FC). (B) Gene ontology analysis for down regulated proteins

**Supplementary Table 1.** NSS scoring. This Excel file reports all the individual NSS scores calculated for each mouse sacrificed as part of this publication.

**Supplementary Table 2.**
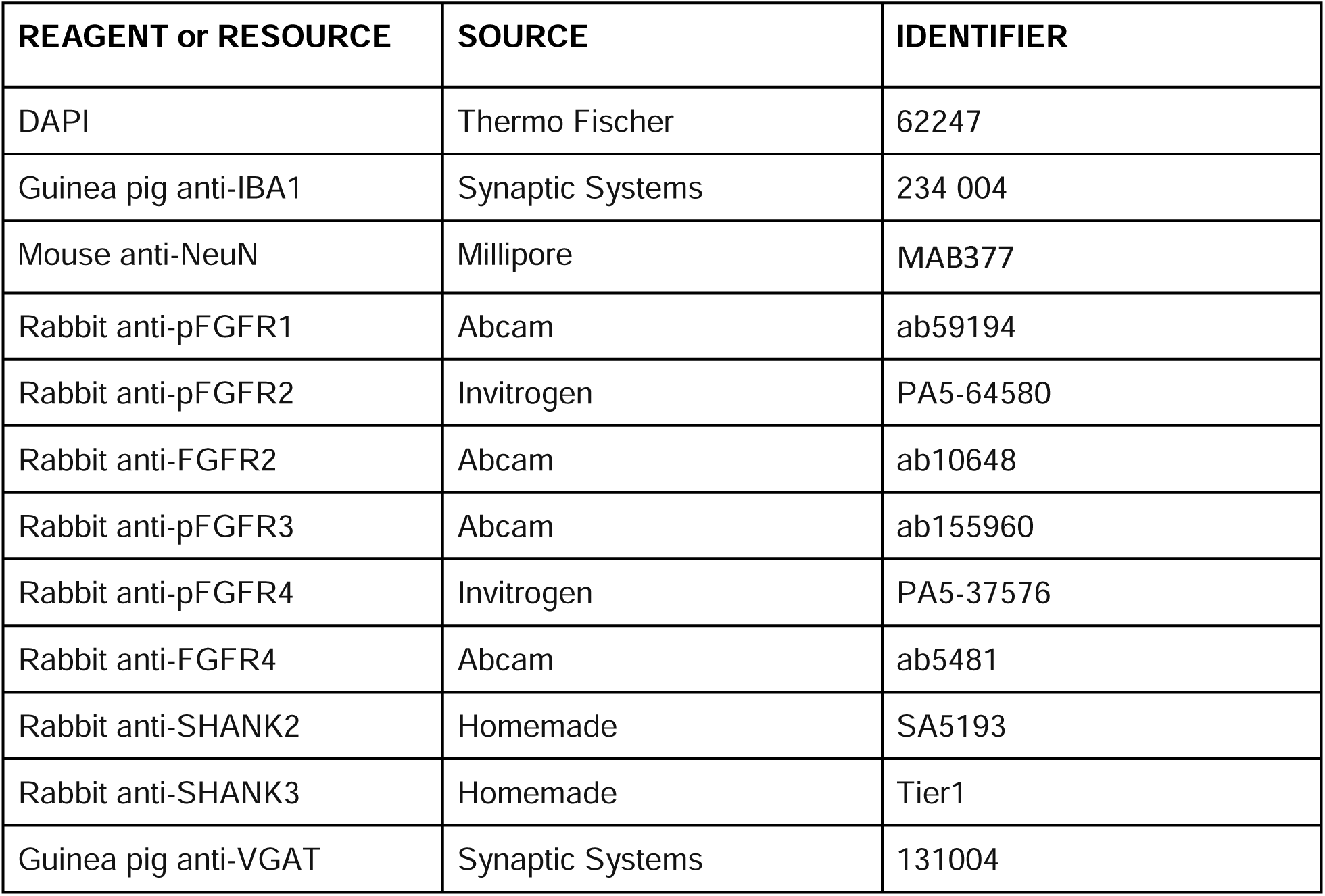
Detailed Antibody list. This list contains all antibody reagents used throughout the publication together with their official name, source and identifier.

## References

Addeo A, Rothschild SI, Holer L, Schneider M, Waibel C, Haefliger S, Mark M, Fernandez E, Mach N, Mauti L, Jermann PM, Alborelli I, Calgua B, Savic-Prince S, Joerger M, Früh M. Fibroblast growth factor receptor (FGFR) inhibitor rogaratinib in patients with advanced pretreated squamous-cell non-small cell lung cancer over-expressing FGFR mRNA: The SAKK 19/18 phase II study. Lung Cancer. 2022 Oct;172:154–159. doi: 10.1016/j.lungcan.2022.08.016.

Bernard C, Exposito-Alonso D, Selten M, Sanalidou S, Hanusz-Godoy A, Aguilera A, Hamid F, Oozeer F, Maeso P, Allison L, Russell M, Fleck RA, Rico B, Marín O. Cortical wiring by synapse type-specific control of local protein synthesis. Science. 2022 Nov 25;378(6622):eabm7466. doi: 10.1126/science.abm7466.

Byrd VM, Kilkenny DM, Dikov MM, Reich MB, Rocheleau JV, Armistead WJ, Thomas JW, Miller GG. Fibroblast growth factor receptor-1 interacts with the T-cell receptor signalling pathway. Immunol Cell Biol. 2003 Dec;81(6):440–50. doi: 10.1046/j.1440-1711.2003.01199.x.

Cangalaya C, Wegmann S, Sun W, Diez L, Gottfried A, Richter K, Stoyanov S, Pakan J, Fischer KD, Dityatev A. Real-time mechanisms of exacerbated synaptic remodeling by microglia in acute models of systemic inflammation and tauopathy. Brain Behav Immun. 2023 May;110:245–259. doi: 10.1016/j.bbi.2023.02.023.

Chandrasekar A, Olde Heuvel F, Wepler M, Rehman R, Palmer A, Catanese A, Linkus B, Ludolph A, Boeckers T, Huber-Lang M, Radermacher P, Roselli F. The Neuroprotective Effect of Ethanol Intoxication in Traumatic Brain Injury Is Associated with the Suppression of ErbB Signaling in Parvalbumin-Positive Interneurons. J Neurotrauma. 2018 Nov 15;35(22):2718–2735. doi: 10.1089/neu.2017.5270.

Chen J, Wang X, Hu J, Du J, Dordoe C, Zhou Q, Huang W, Guo R, Han F, Guo K, Ye S, Lin L, Li X. FGF20 Protected Against BBB Disruption After Traumatic Brain Injury by Upregulating Junction Protein Expression and Inhibiting the Inflammatory Response. Front Pharmacol. 2021 Jan 25;11:590669. doi: 10.3389/fphar.2020.590669.

Dabrowski A, Terauchi A, Strong C, Umemori H. Distinct sets of FGF receptors sculpt excitatory and inhibitory synaptogenesis. Development. 2015 May 15;142(10):1818–30. doi: 10.1242/dev.115568.

Davalos D, Grutzendler J, Yang G, Kim JV, Zuo Y, Jung S, Littman DR, Dustin ML, Gan WB. ATP mediates rapid microglial response to local brain injury in vivo. Nat Neurosci. 2005 Jun;8(6):752–8. doi: 10.1038/nn1472.

Ehlers MD. Activity level controls postsynaptic composition and signaling via the ubiquitin-proteasome system. Nat Neurosci. 2003 Mar;6(3):231–42. doi: 10.1038/nn1013. Erratum in: Nat Neurosci. 2006 Mar;9(3):453.

Farahnak S, McGovern TK, Kim R, O’Sullivan M, Chen B, Lee M, Yoshie H, Wang A, Jang J, Al Heialy S, Lauzon AM, Martin JG. Basic Fibroblast Growth Factor 2 Is a Determinant of CD4 T Cell-Airway Smooth Muscle Cell Communication through Membrane Conduits. J Immunol. 2017 Nov 1;199(9):3086–3093. doi: 10.4049/jimmunol.1700164.

Fröhlich A, Olde Heuvel F, Rehman R, Krishnamurthy SS, Li S, Li Z, Bayer D, Conquest A, Hagenston AM, Ludolph A, Huber-Lang M, Boeckers T, Knöll B, Morganti-Kossmann MC, Bading H, Roselli F. Neuronal nuclear calcium signaling suppression of microglial reactivity is mediated by osteoprotegerin after traumatic brain injury. J Neuroinflammation. 2022 Nov 19;19(1):279. doi: 10.1186/s12974-022-02634-4.

Furusho M, Roulois AJ, Franklin RJ, Bansal R. Fibroblast growth factor signaling in oligodendrocyte-lineage cells facilitates recovery of chronically demyelinated lesions but is redundant in acute lesions. Glia. 2015 Oct;63(10):1714–28. doi: 10.1002/glia.22838.

Grünewald S, Politz O, Bender S, Héroult M, Lustig K, Thuss U, Kneip C, Kopitz C, Zopf D, Collin MP, Boemer U, Ince S, Ellinghaus P, Mumberg D, Hess-Stumpp H, Ziegelbauer K. Rogaratinib: A potent and selective pan-FGFR inhibitor with broad antitumor activity in FGFR-overexpressing preclinical cancer models. Int J Cancer. 2019 Sep 1;145(5):1346–1357. doi: 10.1002/ijc.32224.

Haddad LE, Khzam LB, Hajjar F, Merhi Y, Sirois MG. Characterization of FGF receptor expression in human neutrophils and their contribution to chemotaxis. Am J Physiol Cell Physiol. 2011 Nov;301(5):C1036–45. doi: 10.1152/ajpcell.00215.2011.

Han X, Quarles LD. Multiple faces of fibroblast growth factor-23. Curr Opin Nephrol Hypertens. 2016 Jul;25(4):333–42. doi: 10.1097/MNH.0000000000000240.

Huang Y, Wang F, Li H, Xu S, Xu W, Pan X, Hu Y, Mao L, Qian S, Pan J. Inhibition of Fibroblast Growth Factor Receptor by AZD4547 Protects Against Inflammation in Septic Mice. Inflammation. 2019 Dec;42(6):1957–1967. doi: 10.1007/s10753-019-01056-4.

Jamjoom AAB, Rhodes J, Andrews PJD, Grant SGN. The synapse in traumatic brain injury. Brain. 2021 Feb 12;144(1):18–31. doi: 10.1093/brain/awaa321.

Jassam YN, Izzy S, Whalen M, McGavern DB, El Khoury J. Neuroimmunology of Traumatic Brain Injury: Time for a Paradigm Shift. Neuron. 2017 Sep 13;95(6):1246–1265. doi: 10.1016/j.neuron.2017.07.010.

Ju S, Xu C, Wang G, Zhang L. VEGF-C Induces Alternative Activation of Microglia to Promote Recovery from Traumatic Brain Injury. J Alzheimers Dis. 2019;68(4):1687–1697. doi: 10.3233/JAD-190063.

Kamali S, Rajendran R, Stadelmann C, Karnati S, Rajendran V, Giraldo-Velasquez M, Berghoff M. Oligodendrocyte-specific deletion of FGFR2 ameliorates MOG_35-55_ -induced EAE through ERK and Akt signalling. Brain Pathol. 2021 Mar;31(2):297–311. doi: 10.1111/bpa.12916.

Kerrisk Campbell M, Sheng M. USP8 Deubiquitinates SHANK3 to Control Synapse Density and SHANK3 Activity-Dependent Protein Levels. J Neurosci. 2018 Jun 6;38(23):5289–5301. doi: 10.1523/JNEUROSCI.3305-17.2018.

Klimaschewski L, Claus P. Fibroblast Growth Factor Signalling in the Diseased Nervous System. Mol Neurobiol. 2021 Aug;58(8):3884–3902. doi: 10.1007/s12035-021-02367-0.

Krasemann S, Madore C, Cialic R, Baufeld C, Calcagno N, El Fatimy R, Beckers L, O’Loughlin E, Xu Y, Fanek Z, Greco DJ, Smith ST, Tweet G, Humulock Z, Zrzavy T, Conde-Sanroman P, Gacias M, Weng Z, Chen H, Tjon E, Mazaheri F, Hartmann K, Madi A, Ulrich JD, Glatzel M, Worthmann A, Heeren J, Budnik B, Lemere C, Ikezu T, Heppner FL, Litvak V, Holtzman DM, Lassmann H, Weiner HL, Ochando J, Haass C, Butovsky O. The TREM2-APOE Pathway Drives the Transcriptional Phenotype of Dysfunctional Microglia in Neurodegenerative Diseases. Immunity. 2017 Sep 19;47(3):566–581.e9. doi: 10.1016/j.immuni.2017.08.008.

Li AJ, Suzuki S, Suzuki M, Mizukoshi E, Imamura T. Fibroblast growth factor-2 increases functional excitatory synapses on hippocampal neurons. Eur J Neurosci. 2002 Oct;16(7):1313–24. doi: 10.1046/j.1460-9568.2002.02193.x.

Loane DJ, Kumar A. Microglia in the TBI brain: The good, the bad, and the dysregulated. Exp Neurol. 2016 Jan;275 Pt 3(0 3):316–327. doi: 10.1016/j.expneurol.2015.08.018.

Luo B, Liu Z, Lin D, Chen W, Ren D, Yu Z, Xiong M, Zhao C, Fei E, Li B. ErbB4 promotes inhibitory synapse formation by cell adhesion, independent of its kinase activity. Transl Psychiatry. 2021 Jun 29;11(1):361. doi: 10.1038/s41398-021-01485-6.

Morganti-Kossmann MC, Semple BD, Hellewell SC, Bye N, Ziebell JM. The complexity of neuroinflammation consequent to traumatic brain injury: from research evidence to potential treatments. Acta Neuropathol. 2019 May;137(5):731–755. doi: 10.1007/s00401-018-1944-6.

Parthasarathy G, Pattison MB, Midkiff CC. The FGF/FGFR system in the microglial neuroinflammation with Borrelia burgdorferi: likely intersectionality with other neurological conditions. J Neuroinflammation. 2023 Jan 17;20(1):10. doi: 10.1186/s12974-022-02681-x.

Pottier C, Fresnais M, Gilon M, Jérusalem G, Longuespée R, Sounni NE. Tyrosine Kinase Inhibitors in Cancer: Breakthrough and Challenges of Targeted Therapy. Cancers (Basel). 2020 Mar 20;12(3):731. doi: 10.3390/cancers12030731.

Rajendran R, Rajendran V, Böttiger G, Stadelmann C, Shirvanchi K, von Au L, Bhushan S, Wallendszus N, Schunin D, Westbrock V, Liebisch G, Ergün S, Karnati S, Berghoff M. The small molecule fibroblast growth factor receptor inhibitor infigratinib exerts anti-inflammatory effects and remyelination in a model of multiple sclerosis. Br J Pharmacol. 2023 Jul 3. doi: 10.1111/bph.16186.

Rajendran R, Giraldo-Velásquez M, Stadelmann C, Berghoff M. Oligodendroglial fibroblast growth factor receptor 1 gene targeting protects mice from experimental autoimmune encephalomyelitis through ERK/AKT phosphorylation. Brain Pathol. 2018 Mar;28(2):212–224. doi: 10.1111/bpa.12487.

Rajendran R, Böttiger G, Stadelmann C, Karnati S, Berghoff M. FGF/FGFR Pathways in Multiple Sclerosis and in Its Disease Models. Cells. 2021 Apr 13;10(4):884. doi: 10.3390/cells10040884.

Rehman R, Miller M, Krishnamurthy SS, Kjell J, Elsayed L, Hauck SM, Olde Heuvel F, Conquest A, Chandrasekar A, Ludolph A, Boeckers T, Mulaw MA, Goetz M, Morganti-Kossmann MC, Takeoka A, Roselli F. Met/HGFR triggers detrimental reactive microglia in TBI. Cell Rep. 2022 Dec 27;41(13):111867. doi: 10.1016/j.celrep.2022.111867.

Reinecke M, Brear P, Vornholz L, Berger BT, Seefried F, Wilhelm S, Samaras P, Gyenis L, Litchfield DW, Médard G, Müller S, Ruland J, Hyvönen M, Wilhelm M, Kuster B. Chemical proteomics reveals the target landscape of 1,000 kinase inhibitors. Nat Chem Biol. 2023 Oct 30. doi: 10.1038/s41589-023-01459-3.

Ritzel RM, He J, Li Y, Cao T, Khan N, Shim B, Sabirzhanov B, Aubrecht T, Stoica BA, Faden AI, Wu LJ, Wu J. Proton extrusion during oxidative burst in microglia exacerbates pathological acidosis following traumatic brain injury. Glia. 2021 Mar;69(3):746–764. doi: 10.1002/glia.23926.

Ritzel RM, Li Y, Jiao Y, Lei Z, Doran SJ, He J, Shahror RA, Henry RJ, Khan R, Tan C, Liu S, Stoica BA, Faden AI, Szeto G, Loane DJ, Wu J. Brain injury accelerates the onset of a reversible age-related microglial phenotype associated with inflammatory neurodegeneration. Sci Adv. 2023 Mar 10;9(10):eadd1101. doi: 10.1126/sciadv.add1101.

Roselli F, Livrea P, Almeida OF. CDK5 is essential for soluble amyloid β-induced degradation of GKAP and remodeling of the synaptic actin cytoskeleton. PLoS One. 2011;6(7):e23097. doi: 10.1371/journal.pone.0023097.

Roskoski R Jr. Properties of FDA-approved small molecule protein kinase inhibitors: A 2023 update. Pharmacol Res. 2023 Jan;187:106552. doi: 10.1016/j.phrs.2022.106552.

Roth TL, Nayak D, Atanasijevic T, Koretsky AP, Latour LL, McGavern DB. Transcranial amelioration of inflammation and cell death after brain injury. Nature. 2014 Jan 9;505(7482):223-8. doi: 10.1038/nature12808.

Russo MV, McGavern DB. Inflammatory neuroprotection following traumatic brain injury. Science. 2016 Aug 19;353(6301):783-5. doi: 10.1126/science.aaf6260.

Somebang K, Rudolph J, Imhof I, Li L, Niemi EC, Shigenaga J, Tran H, Gill TM, Lo I, Zabel BA, Schmajuk G, Wipke BT, Gyoneva S, Jandreski L, Craft M, Benedetto G, Plowey ED, Charo I, Campbell J, Ye CJ, Panter SS, Nakamura MC, Eckalbar W, Hsieh CL. CCR2 deficiency alters activation of microglia subsets in traumatic brain injury. Cell Rep. 2021 Sep 21;36(12):109727. doi: 10.1016/j.celrep.2021.109727.

Sternberg CN, Petrylak DP, Bellmunt J, Nishiyama H, Necchi A, Gurney H, Lee JL, van der Heijden MS, Rosenbaum E, Penel N, Pang ST, Li JR, García Del Muro X, Joly F, Pápai Z, Bao W, Ellinghaus P, Lu C, Sierecki M, Coppieters S, Nakajima K, Ishida TC, Quinn DI. FORT-1: Phase II/III Study of Rogaratinib Versus Chemotherapy in Patients With Locally Advanced or Metastatic Urothelial Carcinoma Selected Based on *FGFR1*/*3* mRNA Expression. J Clin Oncol. 2023 Jan 20;41(3):629–639. doi: 10.1200/JCO.21.02303.

Subbiah V, Verstovsek S. Clinical development and management of adverse events associated with FGFR inhibitors. Cell Rep Med. 2023 Oct 17;4(10):101204. doi: 10.1016/j.xcrm.2023.101204.

Tzioras M, Daniels MJD, Davies C, Baxter P, King D, McKay S, Varga B, Popovic K, Hernandez M, Stevenson AJ, Barrington J, Drinkwater E, Borella J, Holloway RK, Tulloch J, Moss J, Latta C, Kandasamy J, Sokol D, Smith C, Miron VE, Káradóttir RT, Hardingham GE, Henstridge CM, Brennan PM, McColl BW, Spires-Jones TL. Human astrocytes and microglia show augmented ingestion of synapses in Alzheimer’s disease via MFG-E8. Cell Rep Med. 2023 Sep 19;4(9):101175. doi: 10.1016/j.xcrm.2023.101175.

Yang JJ, Bertolesi GE, Hehr CL, Johnston J, McFarlane S. Fibroblast growth factor receptor 1 signaling transcriptionally regulates the axon guidance cue slit1. Cell Mol Life Sci. 2018 Oct;75(19):3649–3661. doi: 10.1007/s00018-018-2824-x.

Wang J, Ma MW, Dhandapani KM, Brann DW. Regulatory role of NADPH oxidase 2 in the polarization dynamics and neurotoxicity of microglia/macrophages after traumatic brain injury. Free Radic Biol Med. 2017 Dec;113:119–131. doi: 10.1016/j.freeradbiomed.2017.09.017.

Wang D, Liu F, Zhu L, Lin P, Han F, Wang X, Tan X, Lin L, Xiong Y. FGF21 alleviates neuroinflammation following ischemic stroke by modulating the temporal and spatial dynamics of microglia/macrophages. J Neuroinflammation. 2020 Aug 31;17(1):257. doi: 10.1186/s12974-020-01921-2.

Wang Y, Wernersbach I, Strehle J, Li S, Appel D, Klein M, Ritter K, Hummel R, Tegeder I, Schäfer MKE. Early posttraumatic CSF1R inhibition via PLX3397 leads to time- and sex-dependent effects on inflammation and neuronal maintenance after traumatic brain injury in mice. Brain Behav Immun. 2022 Nov;106:49–66. doi: 10.1016/j.bbi.2022.07.164.

Weaver A, Bossaer JB. Fibroblast growth factor receptor (FGFR) inhibitors: A review of a novel therapeutic class. J Oncol Pharm Pract. 2021 Apr;27(3):702–710. doi: 10.1177/1078155220983425.

Willis EF, MacDonald KPA, Nguyen QH, Garrido AL, Gillespie ER, Harley SBR, Bartlett PF, Schroder WA, Yates AG, Anthony DC, Rose-John S, Ruitenberg MJ, Vukovic J. Repopulating Microglia Promote Brain Repair in an IL-6-Dependent Manner. Cell. 2020 Mar 5;180(5):833–846.e16. doi: 10.1016/j.cell.2020.02.013.

Wu H, Zheng J, Xu S, Fang Y, Wu Y, Zeng J, Shao A, Shi L, Lu J, Mei S, Wang X, Guo X, Wang Y, Zhao Z, Zhang J. Mer regulates microglial/macrophage M1/M2 polarization and alleviates neuroinflammation following traumatic brain injury. J Neuroinflammation. 2021 Jan 5;18(1):2. doi: 10.1186/s12974-020-02041-7.

Zanier ER, Fumagalli S, Perego C, Pischiutta F, De Simoni MG. Shape descriptors of the “never resting” microglia in three different acute brain injury models in mice. Intensive Care Med Exp. 2015 Dec;3(1):39. doi: 10.1186/s40635-015-0039-0.

Zhou S, Li Y, Zhang Z, Yuan Y. An insight into the TAM system in Alzheimer’s disease. Int Immunopharmacol. 2023 Mar;116:109791. doi: 10.1016/j.intimp.2023.109791.

